# Structural basis of PPARγ-mediated transcriptional repression by the covalent inverse agonist FX-909

**DOI:** 10.1101/2025.05.04.651770

**Authors:** Zane T. Laughlin, Liudmyla Arifova, Paola Munoz-Tello, Xiaoyu Yu, Mithun Nag Karadi Giridhar, Jinhui Dong, Joel M. Harp, Di Zhu, Theodore M. Kamenecka, Douglas J. Kojetin

## Abstract

Hyperactivation of peroxisome proliferator-activated receptor gamma (PPARγ)-mediated transcription promotes tumor growth in urothelial (bladder) cancer, which can be inhibited by pharmacological compounds that repress PPARγ activity. FX-909 is a covalent PPARγ inverse agonist currently in phase 1 clinical trials for advanced solid malignancies including muscle-invasive bladder cancer. Here, we compared the mechanism of action of FX-909 to other covalent inverse agonists including T0070907, originally reported more than 20 years ago and misclassified as an antagonist, and two recently reported improved covalent inverse agonist analogs, SR33068 and BAY-4931. Functional profiling and NMR studies reveal that FX-909 displays improved corepressor-selective inverse agonism and better stabilizes a transcriptionally repressive PPARγ LBD conformation compared to T0070907. The crystal structure of PPARγ LBD cobound to FX-909 and NCoR1 corepressor peptide reveals a repressive conformation shared by other covalent inverse agonists. These findings build on recent studies highlighting the pharmacological significance and clinical relevance of transcriptionally repressive PPARγ inverse agonists.

## INTRODUCTION

Peroxisome proliferator-activated receptor gamma (PPARγ) is a nuclear receptor (NR) transcription factor that regulates gene programs that influence cellular differentiation, metabolism, adipogenesis, and insulin sensitization ^1^. NRs are modular domain transcription factors with an N-terminal disordered activation domain (NTD) containing the activation function-1 (AF-1) region, a central DNA-binding domain (DBD), and a C-terminal ligand-binding domain (LBD) containing the activation function-2 (AF-2) coregulator interaction surface. The general mechanism of NR-mediated gene expression occurs via recruitment of transcriptional corepressor and coactivator protein complexes, which bind to the AF-1 in a ligand-independent manner and the AF-2 surface in a ligand-dependent manner ^2,3^. In the absence of ligand, PPARγ recruits corepressor proteins resulting in repression of gene expression ^4^, which is then activated upon binding an agonist ligand. Agonist binding enhances coactivator recruitment to promoter and enhancer regions of chromatin, remodeling of chromatin, recruitment of other transcriptionally machinery, which results in increased expression of PPARγ-regulated gene programs ^5^.

Aside from the well-known adipogenic and insulin sensitizing functions of PPARγ, recent studies have reported that aberrant PPARγ signaling occurs in luminal muscle-invasive bladder cancer. Genomic activation of PPARγ-mediated transcription in bladder cancer can occur via different mechanisms including focal amplification/overexpression of PPARγ or mutations in PPARγ or RXRα, a NR that forms a heterodimer with PPARγ and contributes to PPARγ-mediated control of gene expression ^6–11^. These findings suggested that targeting PPARγ with pharmacological inverse agonists that repress PPARγ-mediated transcription may hold therapeutic utility in the treatment of bladder cancer where PPARγ signaling is hyperactivated.

In 2002, Tularik reported a compound called T0070907 that was described as a covalent PPARγ antagonist due to its ability to block agonist binding and activation of PPARγ-mediated transcription ^12^. However, more than 20 years later it is now understood that T0070907 is not effective at blocking all ligands from binding to PPARγ ^13–16^. Furthermore, biochemical and cellular functional profiling studies revealed that independent of its ability or inability to block binding of other ligands, T0070907 is a corepressor-selective pharmacological PPARγ inverse agonist that represses PPARγ-mediated transcription ^12,17^. NMR studies revealed that T0070907-bound PPARγ LBD populates two long-lived conformations in solution, one that binds to corepressor peptide with high affinity (repressive conformation) and the other that binds coactivator peptide with high affinity (active conformation). A crystal structure of PPARγ LBD cobound to T0070907 and NCoR1 corepressor revealed a NR LBD structural conformation where the critical structural element called helix 12, which is solvent exposed in the active conformation, adopts a solvent occluded conformation within the orthosteric ligand-binding pocket in the repressive conformation ^18^. Subsequent studies revealed that analogs containing a scaffold similar to T0070907 (2-chloro-5-nitrobenzamide) can be optimized to improve PPARγ inverse agonism ^19–22^. NMR studies have also revealed that improved or more efficacious PPARγ inverse agonism, relative to T0070907 as a parent compound, occurs via stabilization of the repressive PPARγ LBD conformation ^19^.

Flare Therapeutics recently announced the development of a first-in-class PPARγ inverse agonist known as FX-909, 3-(5,7-difluoro-4-???-1, 4-dihydroquinolin-2-yl)-4-(methylsulfonyl)benzonitrile, as a drug candidate for advanced urothelial cancer ^23^. FX-909, which is currently in phase 1 clinical trials ^24,25^, is reported to act as a potent and specific covalent PPARγ inverse agonist that represses the expression of PPARγ-mediated transcription, thereby counteracting the cancerous phenotype of bladder cancer cells resulting in tumor regression in mouse models ^26^. A mechanistic biochemical and structural understanding PPARγ LBD conformational bias, or switching between transcriptionally active and repressive conformation, was reported to be critical in the development of FX-909 ^27^. However, the molecular mechanism of action of FX-909 has yet to be reported. Here, we report cellular, biochemical, and structural biology functional profiling of FX-909 along with head-to-head comparison to T0070907 and other recently reported covalent inverse agonists including BAY-4931 ^21^ and SR33068 ^19^ that display improved activity over T0070907.

## RESULTS AND DISCUSSION

### Functional profiling of FX-909 in biochemical and cellular assays

We assembled a set of PPARγ ligands that included FX-909 as well as other covalent ligands including an antagonist (GW9662) ^28^, three inverse agonists (T0070907, SR33068, BAY-4931) ^12,19,21^ and a reference agonist (rosiglitazone) ^29^ (**Figure 1A**). Of the covalent inverse agonists, T0070907 could be considered a parent compound of the other analogs; it was originally reported in 2002 ^12^ as an antagonist and later ligand profiling studies revealed it is a corepressor-selective inverse agonist ^17,18^. We first characterized the compounds using a transcriptional reporter assay where human HEK293T cells were transfected with a plasmid encoding full-length PPARγ along with a reporter plasmid containing three copies of the PPAR-binding DNA response element upstream of the firefly luciferase gene (3xPPRE-luciferase). FX-909 and the other covalent inverse agonists showed a concentration-dependent decrease in PPARγ-mediated transcription with a similar potency (IC_50_ = ∼ 1 nM), whereas GW9662 showed no change in activity and rosiglitazone increased transcription (**Figure 1B**).

**Figure 1.**
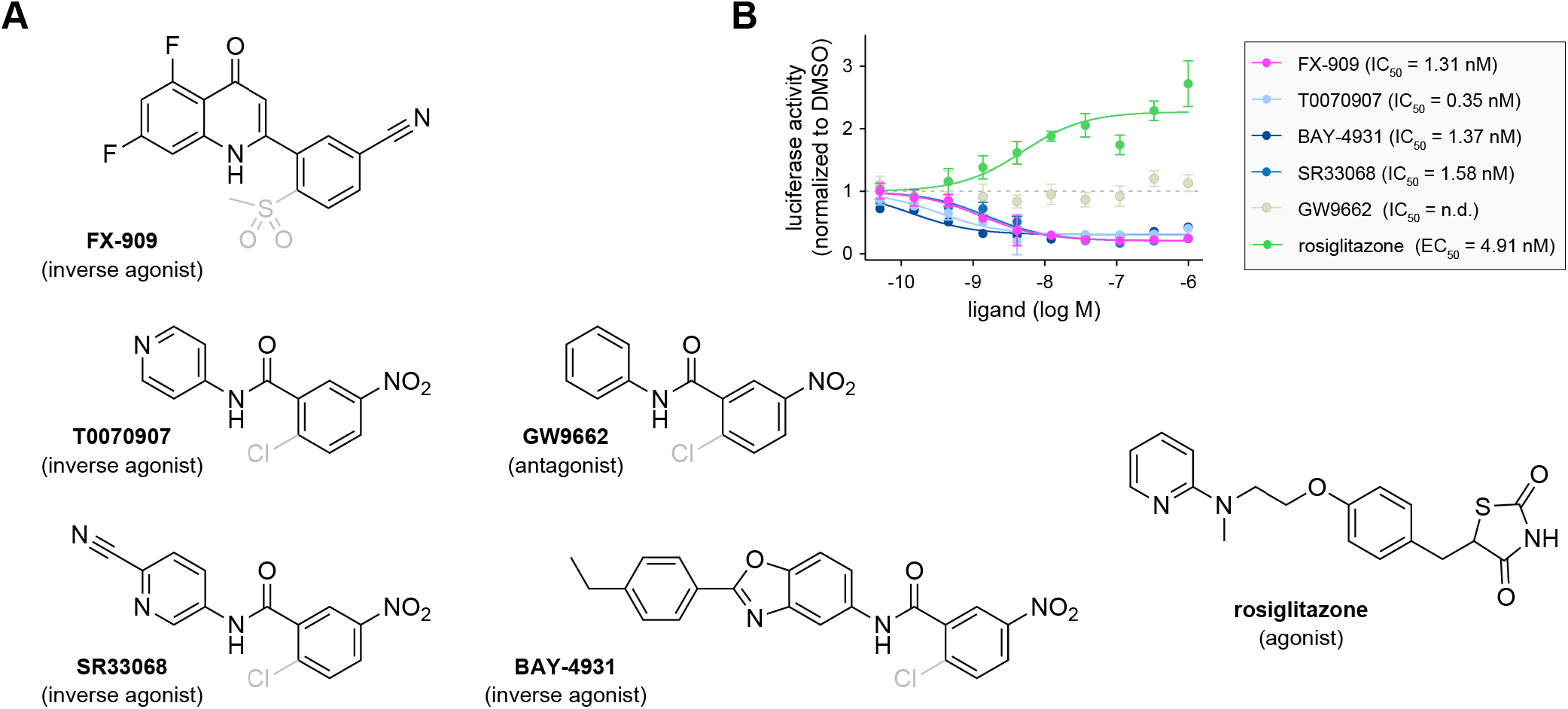
Pharmacological PPARγ ligand set used in this study. (**A**) Chemical structures and pharmacological properties of the ligands. The halogen exchange reaction covalent leaving groups present in all compounds except rosiglitazone are shown in gray. (**B**) Transcriptional reporter assay performed in HEK293T cells transfected with full-length PPARγ expression plasmid and a 3xPPRE-luciferase reporter plasmid. Data were normalized to cells treated with DMSO control, represent mean ± s.d. (n=4), and were fit to a three-parameter sigmoidal dose-response equation to obtain EC_50_/IC_50_ values.

We next characterized the ligand set using biochemical assays that report on ligand-dependent changes in the interaction between PPARγ LBD and FITC-labeled peptides derived from NR coregulator proteins that contribute to PPARγ-mediated transcription in cells including the TRAP220/MED1 coactivator and NCoR1 corepressor. We used time-resolved fluorescence resonance energy transfer (TR-FRET) assays to determine relative ligand potency (or covalent reactivity) and efficacy (via increased or decreased TR-FRET ratio values). FX-909 and the other covalent inverse agonists showed a concentration-dependent increase in NCoR1 corepressor peptide interaction (**Figure 2A**) and decrease in TRAP220/MED1 coactivator interaction (**Figure 2B**), all with similar potency values on the order of 10-30 nM. GW9662 showed relatively little change in efficacy, whereas the rosiglitazone increased TRAP220/MED1 coactivator interaction and decreased NCoR1 corepressor interaction, a profile opposite to the inverse agonists. Relative to the parent compound T0070907, the other covalent inverse agonists show higher TR-FRET efficacy in recruiting the NCoR1 corepressor peptide. These observations suggested the covalent inverse agonist analogs further strengthen the NCoR1 peptide binding affinity compared to T0070907. We performed fluorescence polarization (FP) assays to directly measure binding affinities between the PPARγ LBD and coregulator peptides. Consistent with the TR-FRET efficacy data, the covalent inverse agonist analogs further strengthened NCoR1 corepressor peptide binding affinity (**Figure 2C**) and weakened TRAP220/MED1 coactivator peptide binding affinity (**Figure 2D**). In contrast, rosiglitazone strengthened TRAP220/MED1 binding affinity and decreased NCoR1 binding affinity.

**Figure 2.**
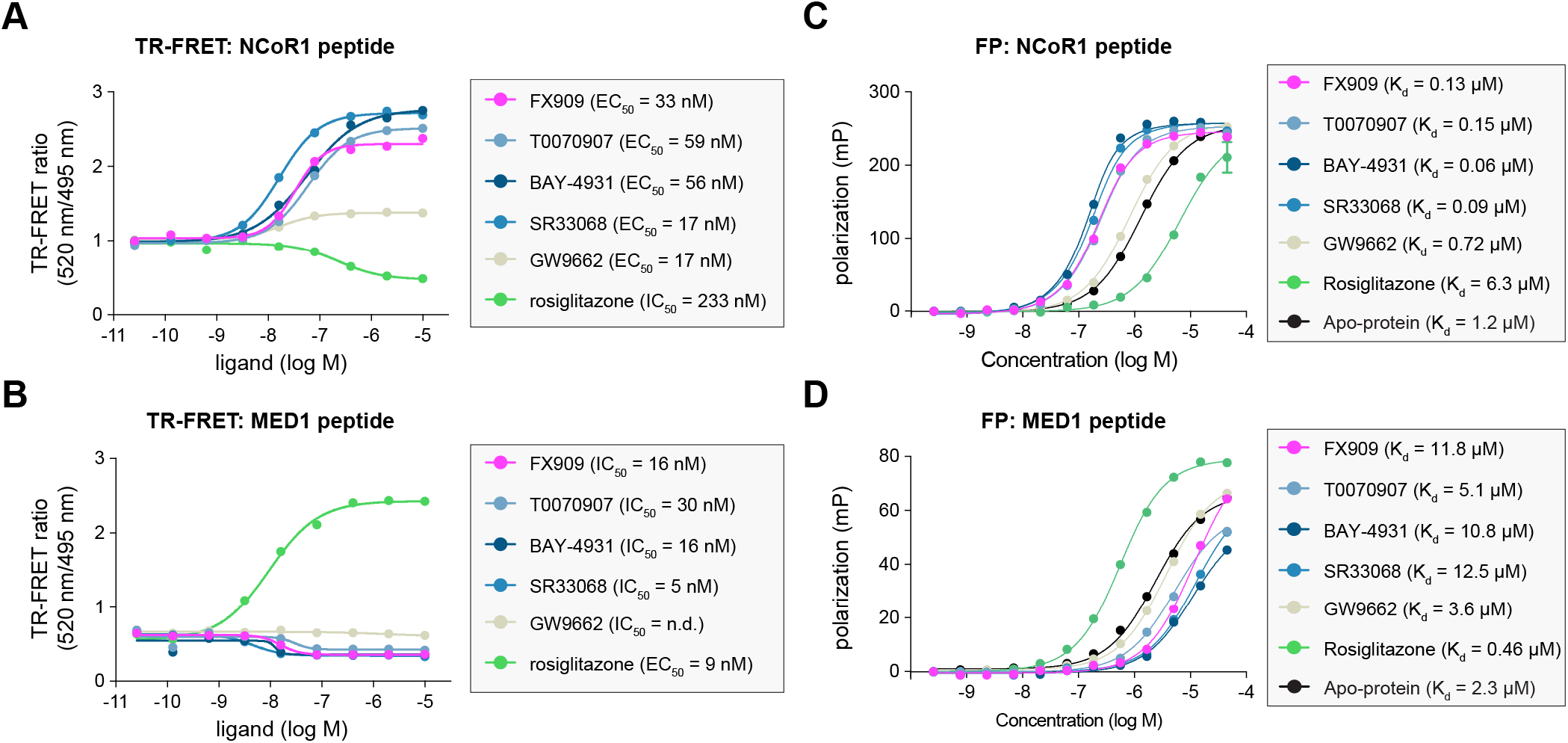
Biochemical coregulator profiling data. (**A**,**B**) Time-resolved fluorescence resonance energy transfer (TR-FRET) coregulator interaction assay performed with (**A**) NCoR1 corepressor peptide or (**B**) TRAP220/MED1 coactivator peptide in the presence of the indicated ligands. Data represent mean ± s.e.m. (n=3) and were fit to a four-parameter sigmoidal dose-response equation to obtain EC_50_/IC_50_ values. (**C**,**D**) Fluorescence polarization (FP) binding assay performed with (**A**) NCoR1 corepressor peptide or (**B**) TRAP220/MED1 coactivator peptide in the presence of the indicated ligands. Data represent mean ± s.e.m. (n=3) and were fit to a quadratic binding equation that assumes binding occurs in a titration binding regime to obtain K_d_ values.

### Crystal structure of PPARγ LBD bound to FX-909 and NCoR1 corepressor peptide

The cellular and biochemical functional profiling data show that FX-909 is a corepressor-selective inverse agonist with improved activity over T0070907 with potency and efficacy values similar to the other improved covalent inverse agonist analogs SR33068 and BAY-4931. To gain insight into the structural basis of FX-909 inverse agonism, we incubated PPARγ LBD with FX-909 followed by addition of NCoR1 corepressor peptide and crystallized the complex, which represents a transcriptionally repressive conformation. We solved a crystal structure of PPARγ LBD bound to FX-909 and NCoR1 corepressor peptide to 2.1Å resolution (**Table 1**). We compared this structure to previously published crystal structures of PPARγ LBD in a repressive conformation bound to NCoR1 peptide and T0070907 ^18^, SR33068 ^19^, BAY-4931 ^21^, and GW9662 ^19^; and in a transcriptionally active conformation bound to TRAP220/MED1 peptide and rosiglitazone ^18^ (**Figure 3**).

**Table 1.**
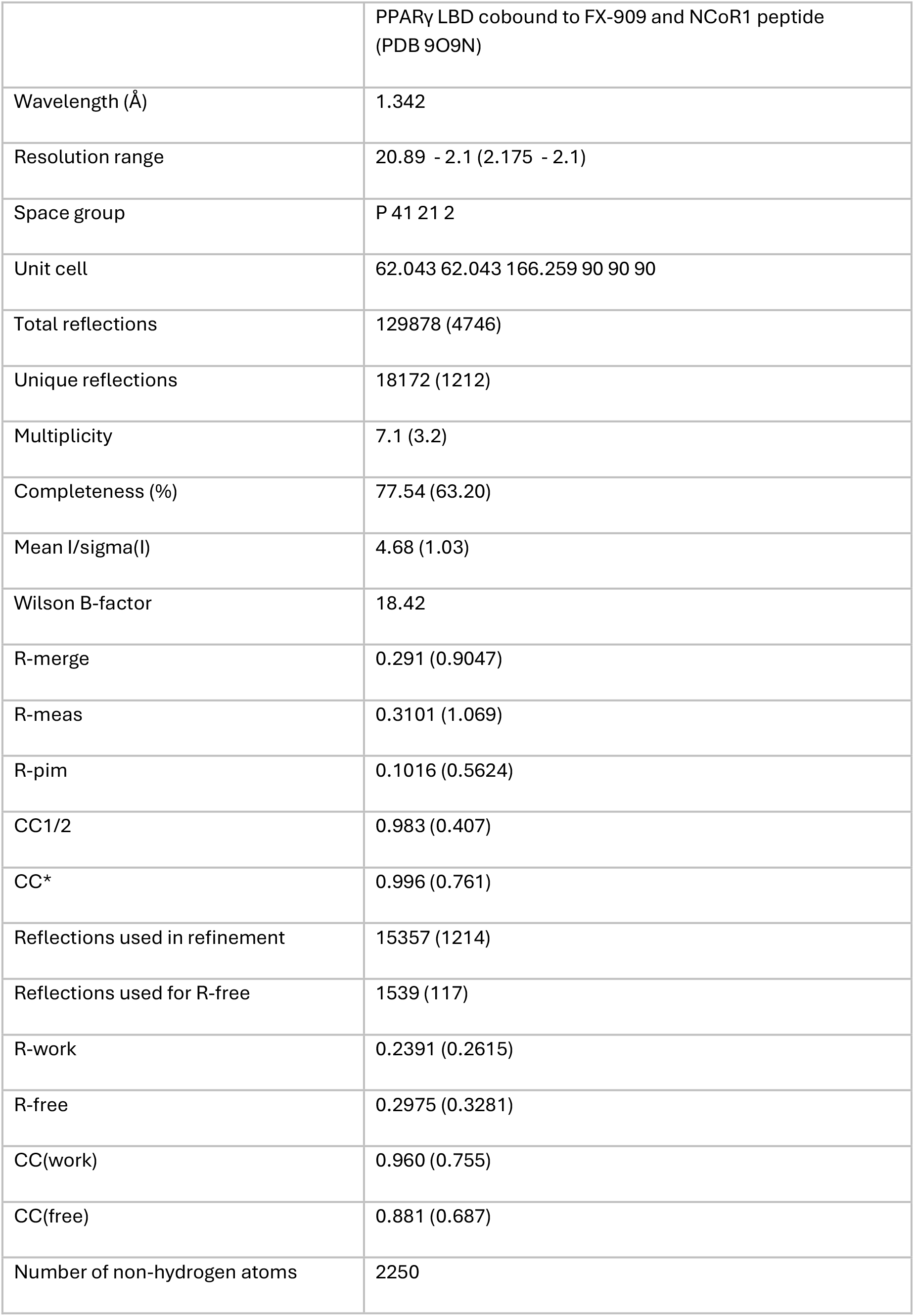
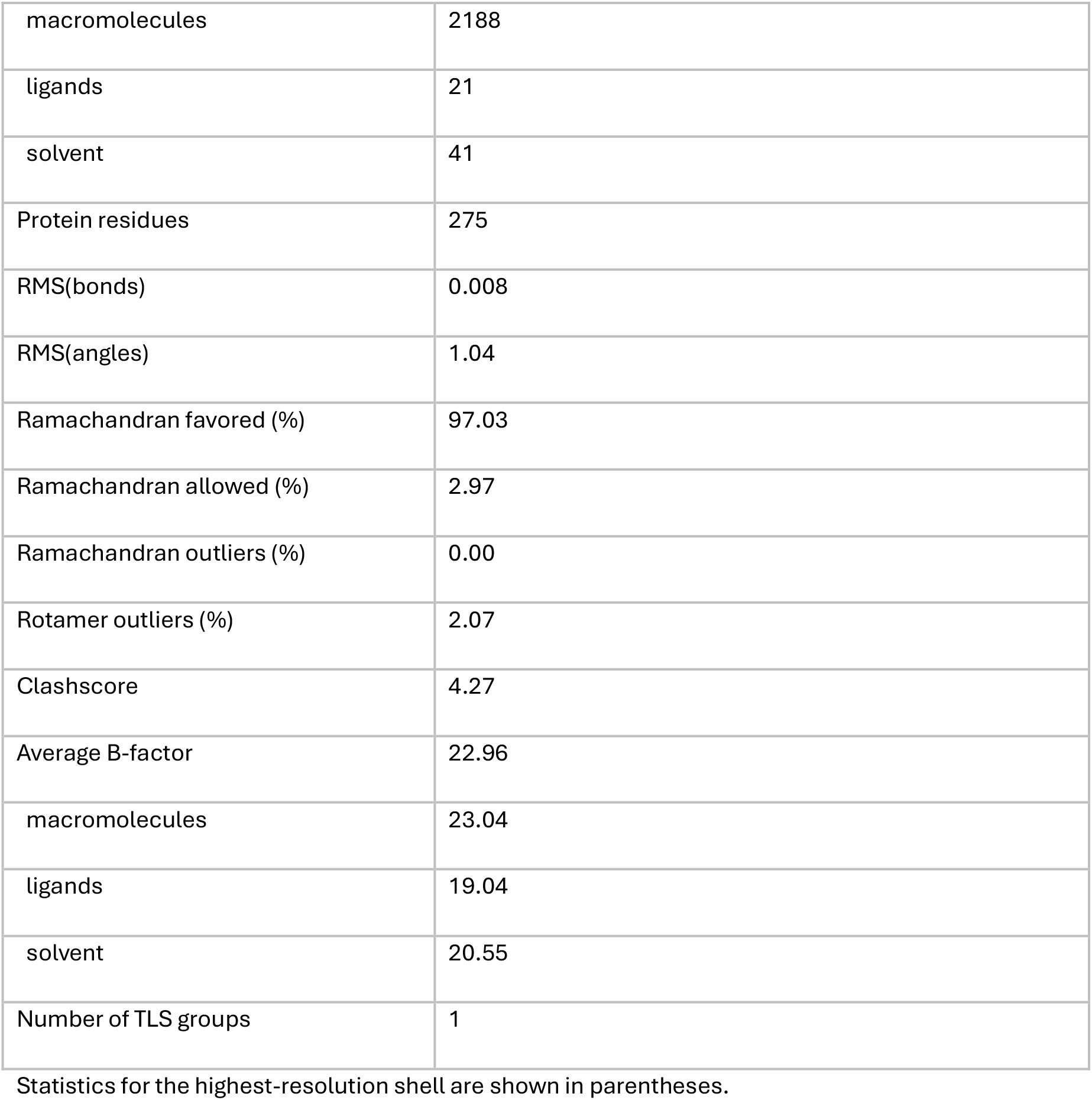
Crystallography data collection and refinement statistics.

**Figure 3.**
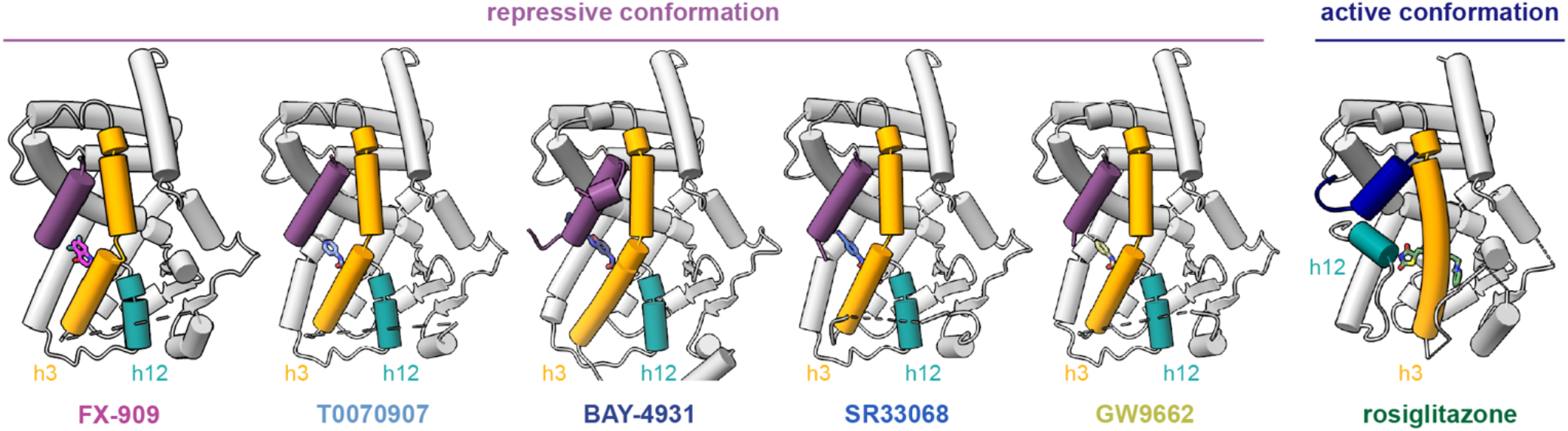
Crystal structures of PPARγ LBD in transcriptionally repressive and active conformations. The repressive conformations crystal structures are bound to NCoR1 corepressor peptide (purple) and covalent ligands including FX-909 (PDB 9O9N), T0070907 (PDB 6ONI), BAY-4931 (PDB 8AQN), SR33068 (PDB 8FKD), and GW9662 (PDB 8FHE). The active conformation crystal structure is bound to TRAP220/MED1 coactivator peptide (dark blue) and rosiglitazone (PDB 6ONJ). The difference between the repressive and active states is most notable in the conformation of helix 3 (orange) and helix 12 (cyan). Ligands are colored according to their annotated names displayed below the structures.

In the active conformation, rosiglitazone is bound within the orthosteric ligand-binding pocket. The thiazolidinedione (TZD) head group forms a hydrogen bond with the hydroxyl-containing side chain of PPARγ LBD residue Y473, which stabilizes the critical structural element helix 12 in a solvent exposed conformation. This solvent exposed helix 12 conformation nucleates an activation function-2 (AF-2) coregulator binding surface conformation that enables high affinity binding of TRAP220/MED1 coactivator peptide containing an LXXLL motif via interactions with two charge clamp residues—K301 on helix 3 and E471 on helix 12. In contrast, the repressive conformation crystal structures adopt a different conformation, where the critical structural element helix 12 adopts a conformation within the orthosteric ligand-binding pocket. This solvent occluded helix 12 conformation leaves the activation function-2 (AF-2) coregulator binding surface completely exposed to enable binding of the NCoR1 corepressor peptide, which is about one helical turn longer than the TRAP220/MED1 coactivator LXXLL motif peptide and would therefore structurally clash if helix 12 adopted a solvent exposed active conformation.

### Covalent ligand interactions with PPARγ

The covalent ligands bind to PPARγ via a halogen exchange mechanism resulting in a covalent bond formed between the substituted phenyl group and the thiol group of residue C285, which points into and positions the ligands within the orthosteric ligand-binding pocket. The FX-909 nitrile group off the substituted phenyl group, near the site of covalent modification with C285, makes a non-covalent polar interaction with the terminal amine side chain of residue K367 (3.04Å) and a weaker interaction (3.72Å) to the thioester side chain of M364 (**Figure 4A**). These interactions also occur with other covalent ligands including T0070907 (**Figure 4B**), BAY-4931 (**Figure 4C**), SR33068 (**Figure 4D**), and GW9662 (**Figure 4E**) that contain nitro groups off the substituted phenyl group, which interact with the M364 thioester more strongly (3.07-3.34Å). In all covalent ligand-bound repressive conformation structures, the amide side chain group of PPARγ residue Q286 makes a polar interaction with a carbonyl group on the ligand, which in FX-909 is off the 5,7-difluoroquinolinone group and in the other ligands comprises the benzamide group. In some of the structures, additional direct and indirect interactions between the Q286 amide side chain are observed with water molecules and nearby residues in PPARγ including H323 and Y477 as well as residue N2260 in the bound NCoR1 peptide. All of the structures show aromatic stacking interactions between the different ligand R-groups — 5,7-difluoroquinolin-4(1H)-one (FX-909), pyridin-4-yl (T0070907), 2-(4-ethylphenyl)benzo[*d*]oxazole (BAY-4931), picolinonitrile (SR33068), and phenyl (GW9662) — extended towards the AF-2 coregulator interaction surface and PPARγ residues with polar side chains including H323, H449, and Y477. Furthermore, the R-group extensions make a variety of direct and indirect, water-mediated polar interactions with side chain of PPARγ residue H323, which likely provide further stability to the repressive conformation. In the structure of PPARγ LBD cobound to rosiglitazone and TRAP220/MED1 coactivator peptide, the TZD headgroup forms polar interactions with side chains of the same polar aromatic residues—H323, H449, and Y473. However, in this active conformation the *N*-methyl-*N*-(2-phenoxyethyl)pyridin-2-amine scaffold of rosiglitazone occupies a large portion of the orthosteric ligand-binding pocket cavity, which as described below is the location where helix 12 occupies in the solvent occluded repressive conformation.

**Figure 4.**
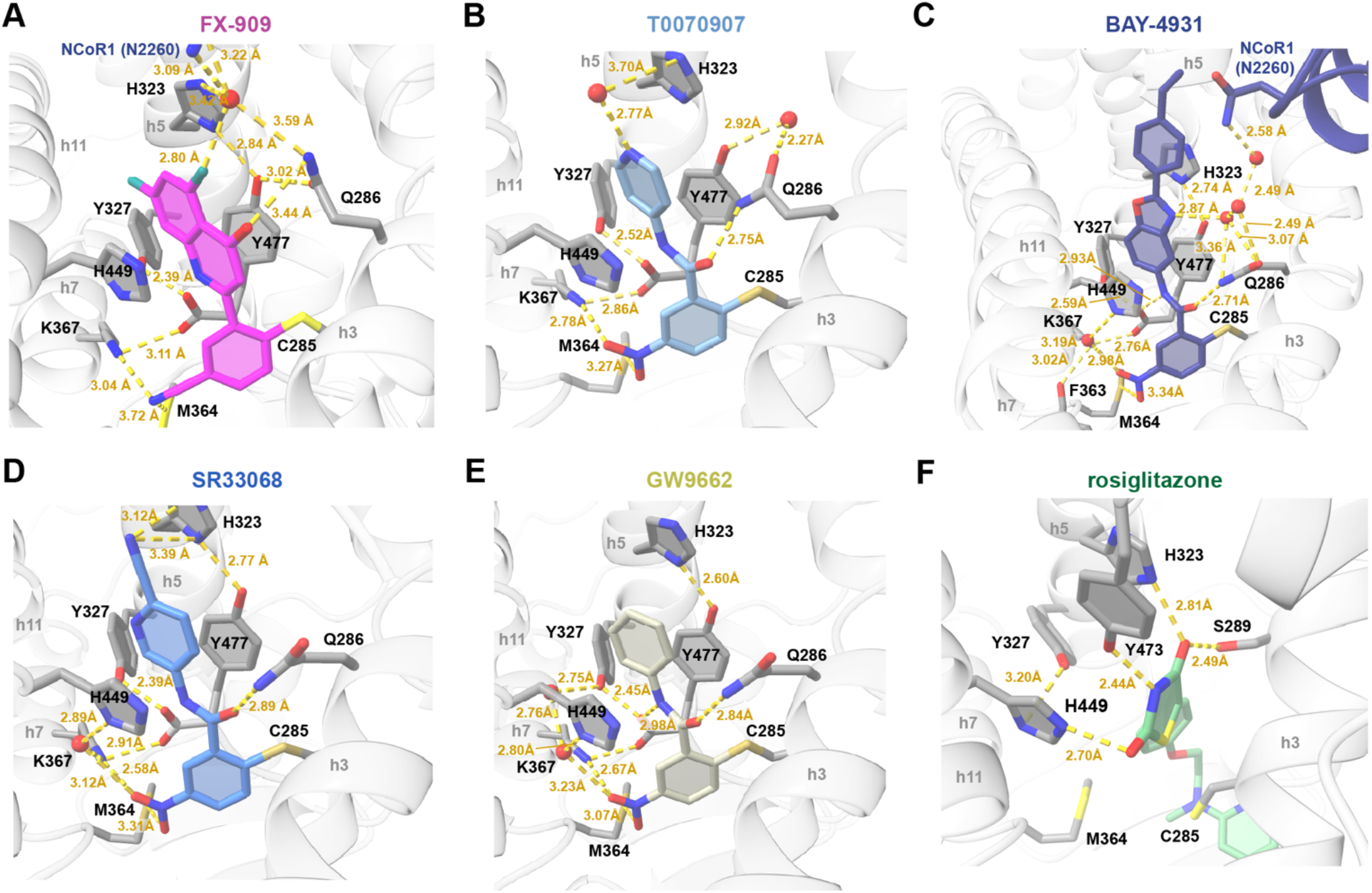
Ligand interactions in the crystal structures. Zoomed in views are shown for PPARγ LBD bound to (**A**) FX-909 (PDB 9O9N), (**B**) T0070907 (PDB 6ONI), (**C**) BAY-4931 (PDB 8AQN), (**D**) SR33068 (PDB 8FKD), (**E**) GW9662 (PDB 8FHE), and (**F**) rosiglitazone (PDB 6ONJ). Secondary structural elements are annotated in gray (e.g., h3, h5, h7, h12). PPARγ LBD residues are annotated in black. Distances are annotated in yellow. Ligands are colored according to the figure panel titles.

### Corepressor peptide interaction at the AF-2 surface

As shown in the TR-FRET and FP data, binding of a covalent inverse agonist ligand to the PPARγ LBD increases the interaction and strengthens binding affinity of NCoR1 corepressor peptide; and decrease and weakens binding affinity of TRAP220/MED1 coactivator peptide. Corepressor and coactivator peptides bind to the PPARγ AF-2 surface through a combination of hydrophobic interactions, where aliphatic residue side chains of the peptide are buried in the AF-2 surface, and polar interactions ^18^. In addition to these hydrophobic interactions, in the FX-909 and NCoR1 peptide cobound crystal structure the peptide makes direct or indirect water-mediated polar interactions with PPARγ residues N286, N294, K301, N312, N314, K319, H323, and Y477 (**Figure 5A**). These residues are also involved in binding NCoR1 peptide when cobound to T0070907 (**Figure 5B**), BAY-4931 (**Figure 5C**), SR33068 (**Figure 5D**), and GW9662 (**Figure 5E**)— although interactions involving N286, H323 and Y477 occur less frequently. In contrast, in the active conformation bound to agonist and coactivator peptide, some of the same residues are involved in the interaction with peptide, most notably K301—one of two charge clamp residues. However, other residues including N312 and K319 make contacts to the N-terminal flexible tail of the TRAP220/MED1 coactivator peptide. Furthermore, several interactions occur between the coactivator peptide and helix 12 in the solvent exposed active conformation including E471—the other charge clamp residue—and I472.

**Figure 5.**
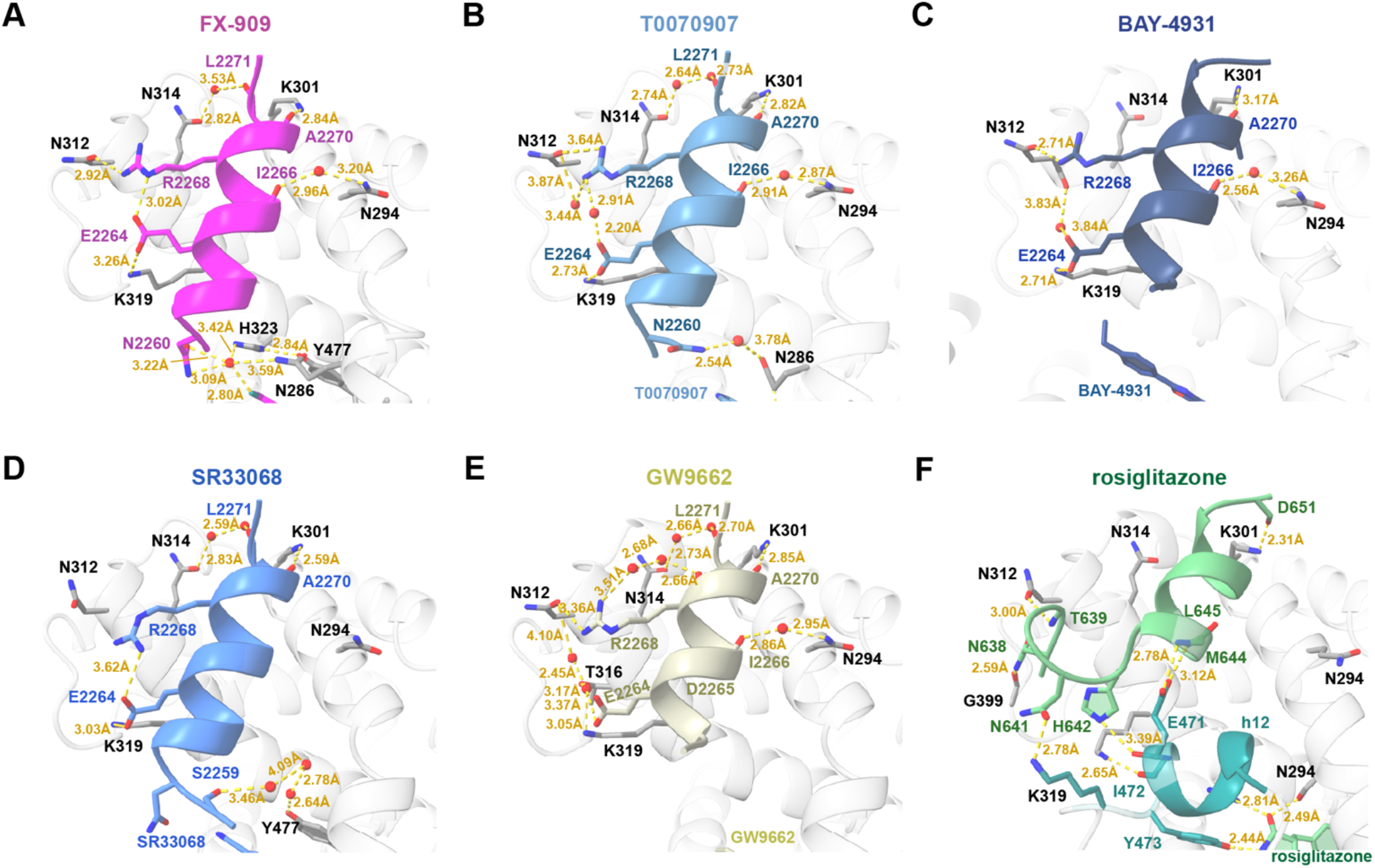
Peptide interactions in the crystal structures. Zoomed in views are shown for PPARγ LBD bound to (**A**) FX-909 (PDB 9O9N), (**B**) T0070907 (PDB 6ONI), (**C**) BAY-4931 (PDB 8AQN), (**D**) SR33068 (PDB 8FKD), (**E**) GW9662 (PDB 8FHE), and (**F**) rosiglitazone (PDB 6ONJ). Secondary structural elements are annotated in gray (e.g., h3, h5, h7, h12). PPARγ LBD residues are annotated in black. Distances are annotated in yellow. Ligands and NCoR1/MED1 peptide residues are colored according to the figure panel titles.

### Repressive helix 12 conformation

In the repressive PPARγ LBD conformation bound to FX-909 and NCoR1 peptide, helix 12 adopts a solvent occluded conformation within the orthosteric binding pocket (**Figure 6A**). Several non-covalent polar interactions likely contribute to stabilizing helix 12 including S342 on β-sheet 2 with D472 on helix 12; R280 on helix 3 with N470 on helix 12; and R288 on helix 3 with E471, D475, and L476 on helix 12. These interactions are conserved in the crystal structures of PPARγ LBD cobound to NCoR1 peptide and T0070907 (**Figure 6B**), BAY-4931 (**Figure 6C**), SR33068 (**Figure 6D**), and GW9662 (**Figure 6E**). Published data show that mutation of R288 (to R288A) and E471 (to E471R) are critical for T0070907-induced recruitment of NCoR1 corepressor peptide and transcriptional repression in cells ^18^. Mutation of R280 (to R280A) also had an effect, though not as pronounced as R280 and E471, which may be explained in the crystals structures as R288 and E471 form a nexus of contacts between helix 12 and helix 3.

**Figure 6.**
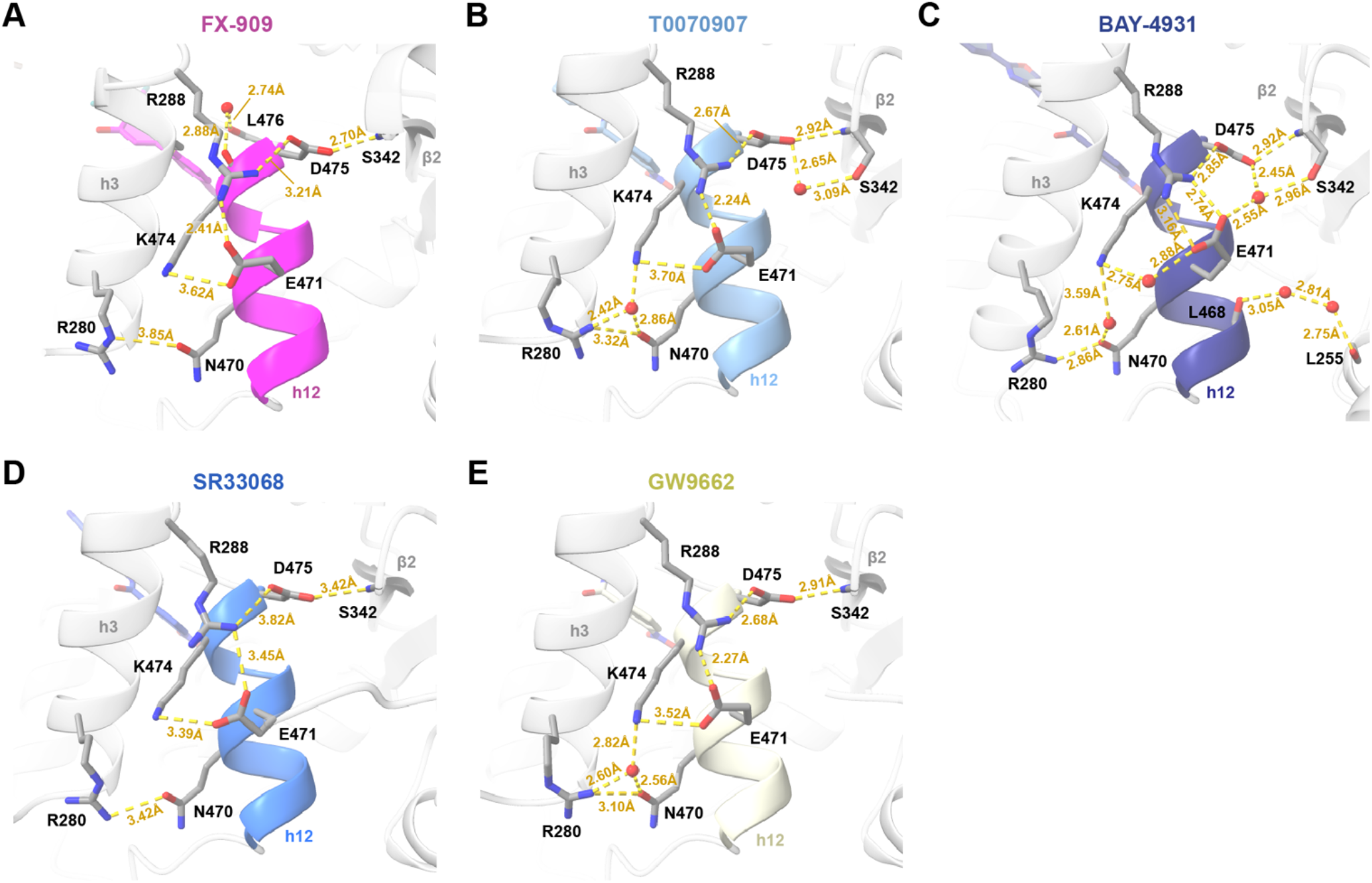
Helix 12 interactions in the repressive conformation crystal structures. Zoomed in views are shown for PPARγ LBD bound to (**A**) FX-909 (PDB 9O9N), (**B**) T0070907 (PDB 6ONI), (**C**) BAY-4931 (PDB 8AQN), (**D**) SR33068 (PDB 8FKD), and (**E**) GW9662 (PDB 8FHE). Secondary structural elements are annotated in gray (h3, β2) except for h12 (colored according to panel titles). PPARγ LBD residues are annotated in black. Distances are annotated in yellow. Ligands and helix 12 (h12) are colored according to the figure panel titles.

### NMR reveals the impact of FX-909 on the PPARγ LBD conformational ensemble

The crystal structures show that FX-909 cobinding with NCoR1 corepressor peptide stabilizes a low energy repressive conformation with helix 12 adopting a solvent occluded position in the orthosteric ligand-binding pocket. However, protein NMR revealed that inverse agonists with graded activity—full inverse agonists that induced robust transcriptional repression, or partial inverse agonists that repress transcription but are less efficacious—shift the PPARγ LBD conformation between a ground state and a fully repressive state in proportion to their relative efficacy ^19^. 2D [^1^H,^15^N]-HSQC TROSY data of ^15^N-labeled PPARγ LBD show that the NMR peak corresponding to G399, a residue near the AF-2 surface that is sensitive to the dynamic PPARγ LBD conformational ensemble exchanging between transcriptionally active and repressive conformations in the absence of ligand ^18^.

We collected 2D [^1^H,^15^N]-HSQC TROSY data of ^15^N-labeled PPARγ LBD in the absence or presence of (**Figure 7A**) FX-909, (**Figure 7B**) T0070907, (**Figure 7C**) BAY-4931, and (**Figure 7D**) SR33068. We compared these NMR data to ^15^N-labeled PPARγ LBD in the absence of ligand (apo/ligand-free), bound to a transcriptionally neutral pharmacological antagonist (GW9662), and bound to a transcriptionally active pharmacological agonist (rosiglitazone). When bound to T0070907, the PPARγ LBD exchanges between transcriptionally active and repressive states, each populated about 50% ^18^. Binding of FX-909 shifts the LBD conformational ensemble more towards a repressive state with a small population of the active state is still observed, similar to improved inverse agonists that we previously reported including SR33068 that show colinear shifting of G399 NMR peak(s) along a diagonal between active and repressive states ^19,30^. BAY-4931 further shifts the LBD conformation towards a repressive conformation, but distinct from FX-909 and SR33068 a lowly populated peak is observed near the repressive conformation peak. This could indicate that when bound to BAY-4931 the repressive LBD conformation exchanges with the active conformation, but the kinetics of exchange are on a different NMR time scale (fast) compared to FX-909 and SR33068 (slow).

**Figure 7.**
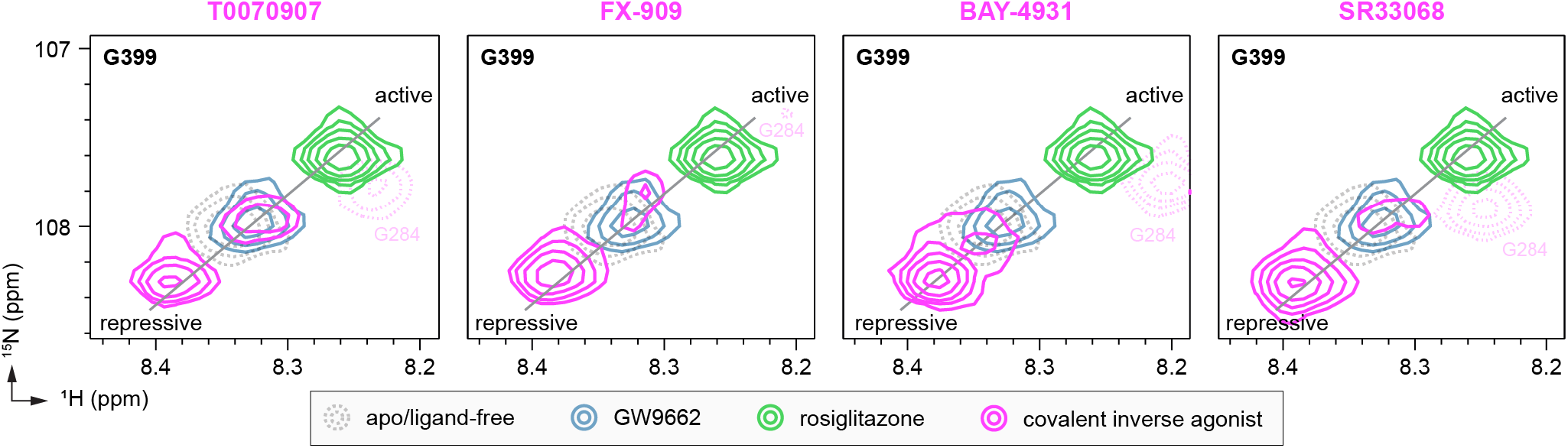
NMR shows that FX-909 shifts the PPARγ LBD conformational ensemble towards a repressive conformation. Overlays of two-dimensional (2D) [^1^H,^15^N]-TROSY-HSQC spectra of ^15^N-labeled PPARγ LBD zoomed into the backbone amide of G399 in the absence or presence of the indicated ligands. The black line denotes the colinear ligand-induced shifting that occurs between repressive and active LBD conformations. The nearby G284 backbone amide NMR peak, which is stabilized by covalent inverse agonists in this region, is colored light pink with a dotted line.

## CONCLUSIONS

If FX-909 is successful in its clinical trials, it will be the first PPARγ-targeted inverse agonist and PPARγ-targeted cancer drug approved as a therapy for use in humans. Our work here shows that, relative to T0070907, FX-909 displays improved function by stabilizing the repressive PPARγ LBD conformation as observed by NMR, which enhances corepressor binding as captured by the crystal structure of PPARγ LBD cobound to FX-909 and NCoR1 peptide resulting in transcriptional repression. The NMR data showing that FX-909 shifts the dynamic PPARγ LBD conformational ensemble from a basal state in the absence of ligand towards a repressive state may explain functional conformational bias that was reported to be critical in the development of FX-909 ^27^. However, compared to BAY-4931 and SR33068, which also shift the LBD ensemble towards a repressive state, extensive medicinal chemistry from Flare Therapeutics likely contributed to the development of a covalent PPARγ inverse agonist with higher specificity (i.e., covalent reactivity) for PPARγ over other potential cellular targets. Furthermore, it is likely that the development strategy resulting in the report of FX-909 included optimization for appropriate *in vivo* drug exposure, which would be critical for use in preclinical studies in animal models and clinical trials in human patients. These are findings among others that we await to hear from Flare Therapeutics in future reports.

## EXPERIMENTAL SECTION

### Compounds and peptides

No newly synthesized compounds are reported this study. Compounds were obtained from commercial sources including MedChemExpress (BAY-4931, FX-909) and Cayman Chemicals (GW9662, rosiglitazone, T0070907) at purity levels are >95%. SR33068 was originally synthesized in our previous study and validated for identify and >95% purity via ^1^H and ^13^C NMR data in our previous publication ^19^. Peptides derived from human NCoR1 (2256-2278; DPASNLGLEDIIRKALMGSFDDK) and human TRAP220/MED1 (residues 638–656; NTKNHPMLM NLLKDNPAQD) were synthesized by LifeTein with an amidated C-terminus for stability, with or without a N-terminal FITC label and a six-carbon linker (Ahx).

### Protein expression, purification, and characterization

PPARγ ligand-binding domain (LBD) protein, residues 203–477 (isoform 1 numbering) with a TEV-cleavable N-terminal hexa-his-tag, was expressed from a pET46 plasmid in Escherichia coli BL21(DE3) cells using autoinduction ZY media (for unlabeled protein), or using M9 minimal media supplemented with ^15^N-labeled ammonium chloride (for labeled protein used in NMR studies). For ZY growth, cells were grown for 5 h at 37 °C, then 1 h at 30 °C, and an additional 12-18 h at 22 °C then harvested by centrifugation (4000g, 30 min). For M9 growth, cells were grown at 37 °C until a sample of the culture reached an OD600nm of 0.6 and then induced with 1.0 mM isopropyl β-D-thiogalactoside (M9) at an OD600nm of 0.6, grown for an additional 12-18 h at 18 °C, and then harvested by the same centrifugation. Cells were resuspended in a buffer containing 50 mM potassium phosphate (pH 7.4), 500 mM KCl, and 10 mM imidazole supplemented with DNase, lysozyme, pepstatin A, leupeptin, and PMSF and lysed by sonication on ice. Cell lysate was clarified by centrifugation (20,000g, 30 min) and filtration (0.2 µm filter). For His-tagged PPARγ-LBD, the protein was purified using Ni-NTA affinity chromatography followed by size exclusion chromatography (Superdex 75) on an AKTA pure in a buffer containing 20 mM potassium phosphate (pH 7.4), 50 mM KCl, and 0.5 mM EDTA. For NMR and studies requiring cleaved protein, the His-tag was cleaved with TEV protease in a buffer containing 20 mM potassium phosphate (pH 7.4), 200 mM KCl, and 0.5 mM EDTA overnight at 4 °C. The TEV-cleaved protein was reloaded on the Ni-NTA column, the flow through collected, and further purified by size exclusion chromatography (Superdex 75) in a buffer containing 20 mM potassium phosphate (pH 7.4), 50 mM KCl, and 0.5 mM EDTA. Purified samples were stored at –80 °C.

### Cellular transcriptional reporter assay

HEK293T (ATCC #CRL-11268) were cultured were cultured according to ATCC guidelines in Dulbecco’s minimal essential medium (DMEM, Gibco). Cells were grown to 90% confluency in T-75 flasks and then 2 million cells were seeded in a 10-cm cell culture dish for transfection using Opti-MEM (Gibco) with full-length human PPARγ (isoform 2) expression plasmid (1.25 ng), and a luciferase reporter plasmid containing the three copies of the PPAR-binding DNA response element (PPRE) sequence (3xPPRE-luciferase) (1.25 ng). After an 18-h incubation, cells were transferred to white 384-well cell culture plates (Thermo Fisher Scientific) at 10,000 cells/well in 20 μL total volume/well. After a 4 h incubation, cells were treated in quadruplicate with 20 μL of either vehicle control (1.5% DMSO in DMEM), threefold serial dilution of ligands for dose response experiments. After a final 18-h incubation, cells were harvested with 20 μL Britelite Plus (PerkinElmer), and luminescence was measured on a BioTek Synergy Neo multimode plate reader. Data were plotted in GraphPad Prism as luminescence vs. ligand concentration as mean ± s.d (n=4) and fit to a three-parameter sigmoidal dose-response curve equation.

### TR-FRET assay

Time-resolved fluorescence resonance energy transfer (TR-FRET) coregulator peptide interaction assays were performed in low-volume black 384-well plates (Greiner) using 23 μL final well volume. Each well contained 4 nM protein (WT or mutant 6xHis-PPARγ LBD), 1 nM LanthaScreen Elite Tb-anti-His Antibody (ThermoFisher #PV5895), and 400 nM FITC-labeled TRAP220/MED1 and NCoR1 peptide in a buffer containing 20 mM potassium phosphate (pH 7.4), 50 mM KCl, 5 mM TCEP, and 0.005% Tween 20. Ligand stocks were prepared via serial dilution in DMSO, added to wells in triplicate, and plates were read using BioTek Synergy Neo multimode plate reader after incubation for 1 hour at 25 °C. The Tb donor was excited at 340 nm, its emission was measured at 495 nm, and the acceptor FITC emission was measured at 520 nm. Data were plotted using GraphPad Prism as TR-FRET ratio (520 nm/ 495 nm) vs. ligand concentration and fit to four-parameter sigmoidal dose-response equation.

### Fluorescence polarization assay

PPARγ LBD was first incubated 1:1 with ligand for 1 hour at 25°C and then serially diluted (1:3) in a buffer containing 20 mM potassium phosphate (pH 8), 50 mM potassium chloride, 5 mM TCEP, 0.5 mM EDTA, and 0.01% Tween20 and plated with 180 nM FITC-labeled TRAP220/MED1 or NCoR1 peptides in low-volume black 384-well plates (Greiner) in triplicate. Plates were incubated at 25 °C for 2 hours, and fluorescence polarization was measured on a BioTek Synergy Neo multimode plate reader at 485 nm emission and 528 nm excitation wavelengths. Data were plotted using GraphPad Prism as fluorescence polarization signal in millipolarization units vs. protein concentration and fit to a quadratic binding equation that assumes binding occurs in a titration binding regime to obtain K_d_ values (equations 6 and 7 in ^31^), which was necessary for the NCoR1 peptide binding data and therefore also used to fit the TRAP220/MED1 peptide binding data.

### X-ray crystallography and structure refinement

Purified PPARγ LBD (concentrated to 10 mg/mL) was incubated with FX-909 at a 1:3 protein/ligand molar ratio overnight, then incubated with NCoR1 peptide at a 1:3 protein/peptide molar ratio and exchanged into a buffer containing 20 mM potassium phosphate (pH 7.4), 50 mM KCl, and 0.5 mM EDTA to remove DMSO and unbound ligands and peptides. Protein complex crystals were obtained after ∼1 month at sitting-drop vapor diffusion against 500 µL of well solution using 24-well format crystallization plates. The crystallization drops contained 1 μL of protein complex sample mixed with 1 μL of reservoir solution containing 0.1 M MES (pH 6.5), 0.2 M Ammonium Sulfate, and 30% PEG 8000. Crystals were flash-frozen in liquid nitrogen before data collection. Data collection was carried out using a Bruker D8 Venture System within the crystallography facility at Vanderbilt University. Data were processed and scaled with the program PROTEUM5 (Bruker). The structure was solved by molecular replacement using the program Phaser ^32^ implemented in the PHENIX package ^33^ using a previously published crystal structure of PPARγ LBD cobound to T0070907 and NCoR1 peptide (PDB code: 6ONI) ^18^ as the search model. The structure was refined using PHENIX with several cycles of interactive model rebuilding in Coot ^34^. Statistics for the crystal structure of PPARγ LBD cobound to FX-909 and NCoR1 peptide generated by PHENIX ^33^ are found in **Table 1**.

### NMR spectroscopy

Two-dimensional (2D) [^1^H,^15^N]-TROSY-HSQC-NMR data were collected at 298 K on a Bruker Avance AV-III 800 MHz NMR spectrometer equipped with a TCI cryoprobe. NMR samples contained 200 μM ^15^N-labeled PPARγ LBD in a buffer containing 20 mM potassium phosphate (pH 7.4), 50 mM KCl, 0.5 mM EDTA, and 10% D_2_O. Samples were incubated with 1 molar equivalent of ligand overnight at 4 °C prior to NMR data collection. Data were collected using Bruker Topspin (version 3); and processed and analyzed using NMRFx ^35^ using published NMR chemical shift assignments for PPARγ LBD bound to rosiglitazone (BMRB accession code 17975) ^36^ and T0070907 (BMRB accession code 50000) ^18^ that were extrapolated to the data herein using the minimum chemical shift procedure ^37^.

## ACKNOWLEDGEMENTS

This work was supported by the National Institutes of Health (NIH) grant R01DK124870 from the National Institute of Diabetes and Digestive and Kidney Diseases (NIDDK). The Vanderbilt Biomolecular NMR facility is supported in part by grants for NMR instrumentation from the NSF-MRI (0922862), acquisition of a 900 MHz Ultra-High Field NMR spectrometer in 2009; NIH (S10 RR025677) for console upgrades on all biomolecular NMR spectrometers in 2009; NIH (R35GM118089-04S1), NIH supplement for the helium liquefier in 2019; NIH S10OD034276 to replace the 800 MHz spectrometer in 2024, accompanied by Vanderbilt University matching funds. The contents of this publication are solely the responsibility of the authors and do not necessarily represent the official views of NIH.

## DATA AVAILABILITY

Crystal structure of PPARγ LBD bound to FX-909 and NCoR1 corepressor peptide was deposited in the PDB under accession code 9O9N. All data generated, analyzed, or used in this study were previously published (BMRB accession codes 17975 and 50000) or available from the corresponding author on reasonable request.

## AUTHOR CONTRIBUTIONS

Z.T.L, L.A., P.M.-T., X.Y., M.N.K.G., J.D., and J.M.H prepared samples, performed experiments, and analyzed data. D.Z. and T.M.K. provided previously synthesized compounds SR33068. D.J.K. conceived, designed, and supervised the research; acquired funding; and analyzed data. Z.T.L. and D.J.K. wrote the manuscript with input from all authors.

